# Industry-wide surveillance of Marek’s disease virus on commercial poultry farms

**DOI:** 10.1101/075192

**Authors:** David A. Kennedy, Christopher Cairns, Matthew J. Jones, Andrew S. Bell, Rahel M. Salathé, Susan J. Baigent, Venugopal K. Nair, Patricia A. Dunn, Andrew F. Read

## Abstract

Marek’s disease virus is a herpesvirus of chickens that costs the worldwide poultry industry over 1 billion USD annually. Two generations of Marek’s disease vaccines have shown reduced efficacy over the last half century due to evolution of the virus. Understanding where the virus is present may give insight into whether continued reductions in efficacy are likely. We conducted a three-year surveillance study to assess the prevalence of Marek’s disease virus on commercial poultry farms, determine the effect of various factors on virus prevalence, and document virus dynamics in broiler chicken houses over short (weeks) and long (years) timescales. We extracted DNA from dust samples collected from commercial chicken and egg production facilities in Pennsylvania, USA. Quantitative polymerase chain reaction (qPCR) was used to assess wild-type virus detectability and concentration. Using data from 1018 dust samples with Bayesian generalized linear mixed effects models, we determined the factors that correlated with virus prevalence across farms. Maximum likelihood and autocorrelation function estimation on 3727 additional dust samples were used to document and characterize virus concentrations within houses over time. Overall, wild-type virus was detectable at least once on 36 of 104 farms at rates that varied substantially between farms. Virus was detected in 1 of 3 broiler-breeder operations (companies), 4 of 5 broiler operations, and 3 of 5 egg layer operations. Marek’s disease virus detectability differed by production type, bird age, day of the year, operation (company), farm, house, flock, and sample. Operation (company) was the most important factor, accounting for between 12% and 63.4% of the variation in virus detectability. Within individual houses, virus concentration often dropped below detectable levels and reemerged later. These data characterize Marek’s disease virus dynamics, which are potentially important to the evolution of the virus.

## Introduction

Marek’s disease (MD), caused by Marek’s disease virus (MDV, *Gallid herpesvirus II*), is an economically important disease of chickens. Since the development of the first vaccine against this disease, mass vaccination has been a key feature in sustaining industrial-scale poultry production (27). MD vaccines are described as “leaky”, because they protect vaccinated hosts from developing clinical signs of disease, but they nonetheless allow for infection and onward transmission of the virus (23, 38, 47). This means that the virus can persist and potentially evolve in vaccinated flocks (39). Nevertheless, very little is known about the distribution of the virus in the field. Here we surveilled virus across the industry by sampling dust (the infectious vehicle) from commercial chicken facilities located throughout Pennsylvania from 2012 to 2015. We use these data to ask where MDV is found, how its prevalence differs across the industry, and how its concentration changes within flocks over time.

MDV is a herpesvirus (9) that is transmitted through inhalation of virus-contaminated dust (13). Once inside a host, the virus goes through an incubation period of about one week, after which new virus particles are first shed from feather follicle epithelial cells (3, 22). The shedding of this infectious virus co-occurs with the shedding of epithelial cells, and so the virus can be found in “chicken dust” (10), a by-product of chicken farming made up of sloughed off epithelial cells, feathers, fecal material, chicken feed, and bedding material (12). Once shedding is initiated, it is thought to occur for the rest of the chicken’s life (47).

MD was first described over a century ago as a relatively mild polyneuritis condition in chickens. Over time the disease has increased in severity in unprotected chickens due to altered rearing conditions and evolution of the virus (31, 39, 46). Two generations of MD vaccines have been undermined by virus evolution, and this evolutionary trajectory has been well documented (46). Whether the efficacy of existing vaccine control strategies will decline in the future is an open question (28), whose answer partially depends on the ecology of the virus. This is because evolutionary outcomes can vary greatly depending on ecological details, which in this case depend on where in the industry the evolution is occurring (1, 39).

Early efforts to quantify MDV prevalence in the field used serological data to demonstrate that infection was extremely prevalent (5, 11, 20, 47). Clinical disease and production losses coupled with these observations motivated near-universal vaccination in commercial poultry farming in the United States and many other nations. More recently, virus prevalence has been inferred from condemnation data (26, 34, 45) and questionnaires (15), but the reliability of these methods are limited by changes in disease and perception of disease that may occur irrespective of virus dynamics (26). The development of quantitative polymerase chain reaction (qPCR) protocols specific for MDV have made it possible to detect and quantify virus collected from field settings (2, 3, 21). Four studies have used qPCR methods to study field samples to study virus dynamics in Australia (17, 37, 44) and Iraq (43). There are many differences in chicken and egg production between these countries and the United States, perhaps most notably that vaccination is nearly universal among commercial farms in the United States (44). Here we performed quantitative polymerase chain reaction (qPCR) on samples collected from chicken farms throughout Pennsylvania, USA, to directly examine MDV dynamics on commercial poultry farms. The farms used in our study encompass much of the diversity of industrial-scale commercial chicken-meat and egg production.

Commercial poultry farming is highly structured (fig. 1). Industrialized commercial chicken production is broadly divided into egg laying birds, broiler birds, and layer-breeder or broiler-breeder birds Each have potentially different natural histories, genetics, and management practices. Further structure exists within these production types, because of differing management practices between operations (companies), for example from targeting particular sectors of the poultry market (e.g. kosher, organic, live bird market, cage-free eggs, etc.), or by sharing biosecurity practices, equipment, and feed mills. Within an operation the behaviors of the people who manage the birds on the farm could in turn affect virus dynamics. Within single farms, there are usually multiple houses. Within these houses, there are successive flocks of birds. Our goal was to quantify the relative importance of these factors on the variation we observed in the prevalence of MDV. This is a critical first step in evaluating risk factors both for disease outbreaks, and for virus evolution that might undermine current vaccine strategies and lead to increased pathogen virulence.

**Figure 1:**
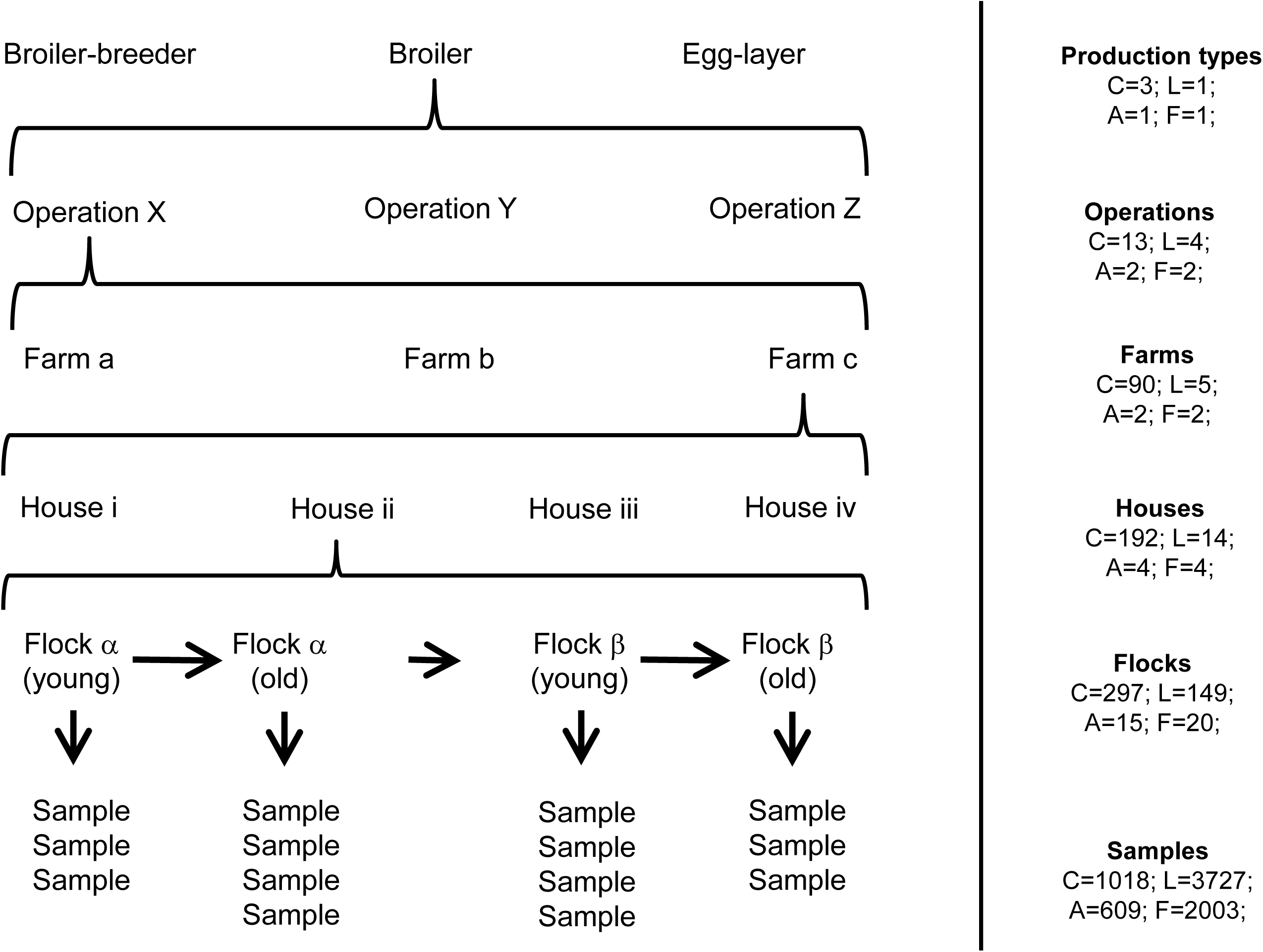
The structure of the data in our study. Left panel: a schematic example of a sampling hierarchy generated by the structure of the poultry industry. Reading from the bottom up, multiple samples were collected from a single flock, multiple flocks were reared in a single house over time, multiple houses were located on a single farm, multiple farms were associated with a single operation (company), and multiple operations were rearing chickens that typically belonged to a single production type. This created a nested hierarchical structure in the data. One example of such a hierarchy is shown here. Right panel: the actual number of unique levels are given by “C” for the cross-sectional data, “L” for the longitudinal data, “A” for the air tube data, and “F” for the feather tip data.

## Methods

### Background

Pennsylvania has commercial scale production of both chicken meat and eggs. Most broiler flocks follow an all-in, all-out approach. Some, however, especially farms rearing colored breeds have multiple ages per premises, while maintaining all-in, all-out practices for individual houses. Down time is typically at least one week, but can range from as little as one day to in excess of several weeks. Most of these farms are cleaned out completely during this down time between flocks, and the farms typically do not re-use litter. Breeder flocks use all-in, all-out approaches for each house with cleaning and disinfecting before new birds are placed. Nevertheless, some have multiple ages on single premises in different houses. Caged layers are typically reared on multi-house complexes, where each house follows an all-in all-out system with cleaning and disinfecting between flocks. Different houses, however, remain populated with different aged birds to achieve continuous egg production. Floor layers are typically reared on premises with one house and one age of bird, or two houses usually of different age from each other. Each house typically follows an all-in, all-out approach with cleaning and disinfection before restocking.

Three live vaccine virus strains are used on Pennsylvania farms to control MD: HVT, SB-1, and Rispens. These strains are related but not identical to wild-type virus. Once vaccinated, a bird can shed these vaccine strains (3, 22), and so we used the primer-probe combination of Baigent et al. (2) that is capable of quantifying wild-type virus in the presence of each of the vaccine strains. This specificity is necessary because almost all chickens in Pennsylvania reared for commercial production are vaccinated against MD. Broiler chickens are typically vaccinated using a combination of HVT and SB-1, although Rispens vaccine virus is used under some circumstances. Egg laying chickens and broiler-breeder chickens are typically given Rispens vaccination, often in combination with HVT and/or SB-1. This was confirmed in our samples through the detection of the Rispens vaccine virus in at least some dust samples from each of these operations. In Supplement A.1, we show that HVT and SB-1 detection in dust is uncorrelated with wild-type virus detection, and that Rispens vaccine virus is negatively correlated with wild-type virus detection.

### Sample collection

The dust that collects on fan covers, or “louvers”, shows less spatial variation in virus concentration than dust that collects on ledge-like surfaces (Supplement A.2), and so samples used in this study were collected by scraping dust from fan louvers. Logistical constraints including those imposed by biosecurity concerns, industry participation, total availability of farms, and time-varying presence of chicken cohorts resulted in a sampling schedule best described formally as haphazard rather than random. Given these constraints, we visited and collected dust from as many different farms as possible to gain insights into whether and where the virus was detectable. A summary of our sample sizes is available in fig. 1. Between two and six samples were collected from each house during each visit. In total, we visited 104 unique commercial combinations of farm and operation (three farms changed operations during surveillance). These combinations were comprised of 29 broiler-breeder facilities, 52 broiler facilities, and 23 egg-laying facilities (no egg-breeder facilities were included). On five broiler farms where high concentrations of virus were detected, we collected at approximately weekly intervals to quantify changes in virus concentration over time (hereafter referred to as the “longitudinal data”). Each of these five farms was visited between 48 and 133 times (mean 98.4). This subset of data includes 3727 samples, collected across 149 flocks, reared in 14 houses on five farms representing 4 operations (fig. 1). We quantified MD prevalence using all fan dust samples with the exception of those from these five farms and 103 other samples for which bird age was unavailable. We refer to this subset of data as the “cross-sectional data.” This subset is comprised of 1018 samples, collected from 297 flocks, reared in 192, located on 90 farms, belonging to 13 operations, with 3 production types (fig. 1). All fan dust samples collected during this study are being stored indefinitely at -80 °C.

On two of the farms included in the longitudinal data study, we also collected data on airborne virus concentration and host infection status. Airborne virus concentration was assessed by securing six 1.5 ml centrifuge tubes to the arms, hips, and legs of two of the authors during routine dust collection. Tubes were oriented horizontally with tops pointing to the front of the collector’s body, opened upon entering the house and closed upon leaving. This period lasted approximately fifteen to twenty minutes. These data are hereafter referred to as the “air tube data.” They are comprised of 609 samples, from 15 flocks, reared in 4 houses, on 2 farms, associated with 2 operations (fig. 1). Both farms reared broiler chickens. Feathers were also collected from individual birds on these same farms as follows. Two feathers were plucked from the breast of each target bird. The pulpy proximal end of each feather was clipped and placed into its own centrifuge tube. Scissors used to clip feathers were cleaned between birds using 70% isopropyl alcohol wipes. Ten total birds were sampled from each house during each visit (hereafter referred to as the “feather tip data”). Target birds for feather collection were chosen such that they were spatially distributed throughout the house. Individual birds were selected at the discretion of the collector with a goal of random selection. To account for the possibility of airborne virus contamination, we also had two control tubes, one that was left open during the collection of a single feather from a single bird, and one that was left open during the collection of feathers from all ten birds. These control tubes are distinct from the air tube samples, which were collected immediately before feather samples. In total, we tested 2003 feathers, from 20 flocks, and 4 houses (fig. 1). Feather sampling was approved by the Institutional Animal Care and Use Committee of The Pennsylvania State University (IACUC Protocol#: 46599)

### qPCR

All samples were brought back to the lab and stored at 4 °C prior to processing. Detailed methods regarding DNA extraction and qPCR can be found in Supplement A.3. Dust samples collected from fans were processed in duplicate using a modified version of the protocol of Baigent et al. (2). Methods were similar for air tube and feather tip samples, but these samples were processed in singlicate. DNA for all samples were captured in a final elution volume of 200 *μ*l, and 4 *μ*l of this undiluted elution were used in each qPCR reaction.

## Statistical analysis

### Analysis of the cross-sectional data

All analyses were performed in the R statistical computing language (36). To study the variation in the presence and absence of MDV across chicken dust samples, we treated our qPCR data as binomial data on a logit scale with those qPCR runs that had at some point crossed the qPCR fluorescence threshold treated as positive outcomes, and those that had not treated as negative outcomes. This method was similar in effect to running a traditional PCR and checking for the amplification of a target using gel electrophoresis. In practice, our limit of detection was approximately 100 template DNA copies per mg of dust (Supplement A.4), which is close to the concentration of virus that would be expected if about 20 to 50 chickens were infected per flock of 30,000 chickens and virus was randomly mixed throughout the dust (Supplement A.5). Feather tip data were similarly treated as binomial data (Supplement A.6).

We analyzed the data using Bayesian generalized linear mixed effects models (7, 16, 18). Justification for the modeling choices below can be found in Supplement A.7. Our analysis was performed using the function ‘MCMCglmm’ (18) with family set to “categorical”, and “slice” sampling. Depth of coverage ranged from 1 to 90 dust samples, with a median of 6 (fig. 2). Models included random effects for “Operation”, “Farm”, “House”, “Flock”, and “Sample” to account for these levels of clustering in the data. For example, including an effect of “Sample” allowed us to distinguish between technical and biological variation in virus detection. For each random effect, we used inverse Wishart priors with scale 5 and degrees of freedom 3 (Supplement A.8). Models also included fixed effects of “Production type”, “Collection date”, and “Bird age”. For each fixed effect, we used univariate normal priors with mean 0 and standard deviation 7 (Supplement A.8). Production type was fit as a categorical factor with levels “broiler”, “broiler-breeder”, and “layer”. Collection date was fit as two continuous factors, one the sine and one the cosine of 2π/365 times the calendar day that a sample was collected, to capture seasonal variation (26). Bird age was fit as a categorical factor using a spline with knots at cohort ages of 21, 42, 100, and 315 days (19). The spline was generated using the ‘bs’ function in the package “splines”. We generated five candidate models consisting of the full model that contains all of the factors listed above, the three models that lacked exactly one of these fixed effects, and one model that lacked the random effect of “Sample”. We explored the importance of the other random effects by examining the magnitude of their estimated effect sizes.

**Figure 2:**
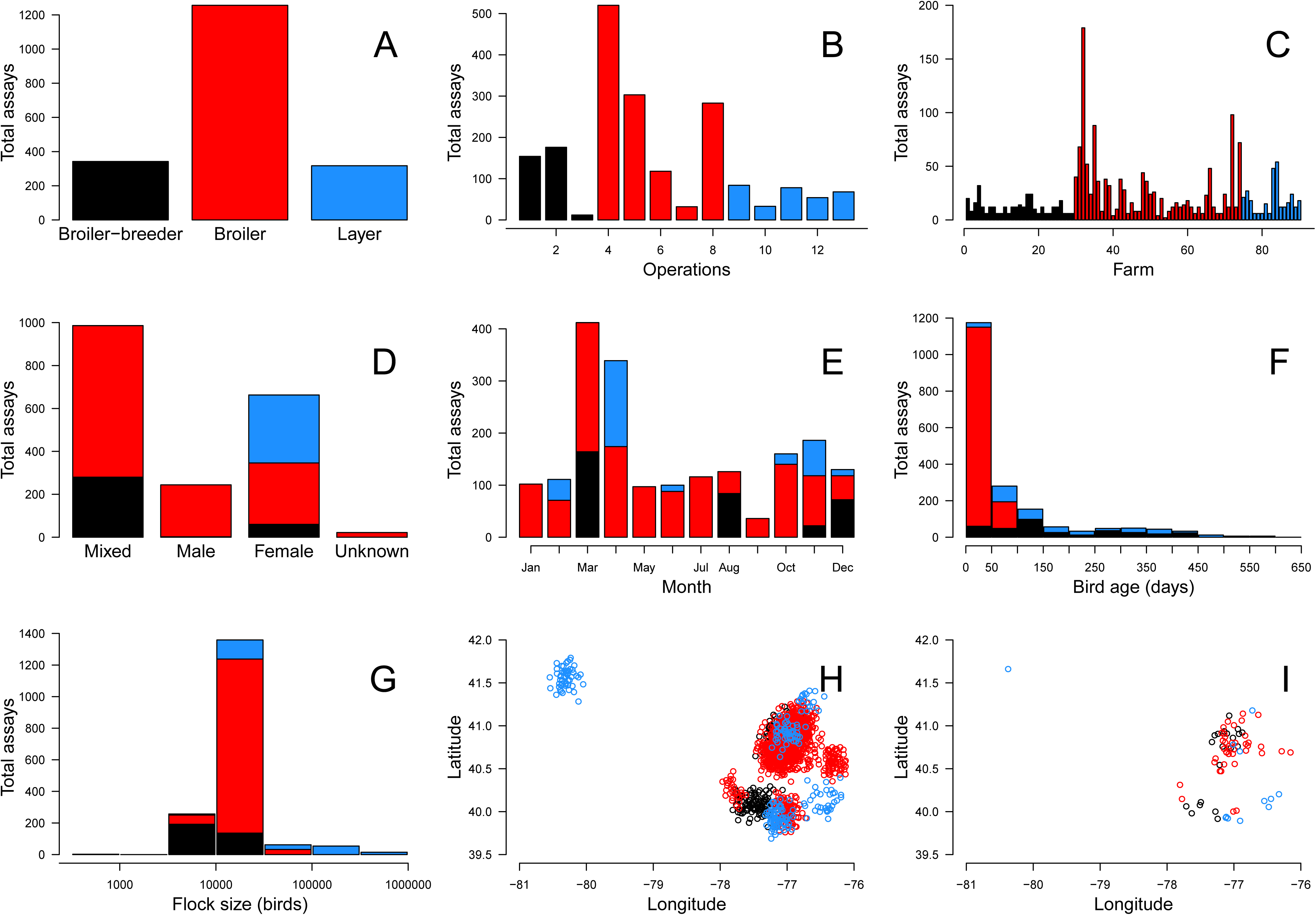
Summary plots of the cross-sectional data depicting the number of assays that were performed as a function of production type (A), operation (B), farm (C), sex (D), month of the year (E), bird age (F), and flock size (G). For example, in panel B, 520 assays were run for samples collected from operation 4. Also depicted are the approximate locations of origin of each sample (H) and each farm (I). Note that to maintain farm location anonymity, normal random variables with mean 0 and standard deviation 0.1 were added to the points when plotting latitude and longitudes in H and I. In all plots, black color depicts breeder facilities, red color depicts broiler facilities, and blue color depicts layer facilities.

We ran each model for 4.1 × 10^6^ iterations with a burn in of 1 × 10^5^ steps, and a thinning interval of 2 × 10^3^. This resulted in 2000 parameter samples for each model run. This process was repeated to generate a total of three chains for each model. Posterior convergence was tested in three steps, following Kennedy et al. (25). The models were then compared using the Deviance Information Criterion (DIC). DIC is a tool, in many ways similar to the Akaike Information Criterion (AIC), that is useful for comparing the relative goodness of fit of various models (42). To foster model comparison, we presented ∆DIC scores, which are the differences in DIC between the best model and each alternative model. Like AIC, there is no precise threshold for significance of ∆DIC scores, but Bolker (6) argued that it is on the same scale as AIC. We therefore followed the suggested rule of thumb for AIC (8) that ∆DIC scores less than 2 suggest substantial support for a model, scores between 3 and 7 indicate considerably less support, and scores greater than 10 suggest that a model is very unlikely.

We also explored the importance of model factors using fraction of variance explained (*R*^2^) where the calculation of *R*^2^ was modified for use with generalized linear mixed models (29). We presented marginal *R*^2^ and conditional *R*^2^ values that describe the fraction of variance on the latent scale of the data that can be attributable to fixed and fixed plus random effects, respectively. We then extended this method to explore the contribution to *R*^2^ that can be attributed to each single factor in the model. Credible intervals for all estimates came from the posterior distributions of the fitted models.

We explored the statistical significance of differences between production types by performing pairwise comparisons on the estimated effect sizes of production type. In practice, this was done by asking what fraction of samples from the posterior estimated a larger effect size for production type level 1 than for production type level 2 or the reverse. This value was multiplied by two to account for it being a two-tailed hypothesis test. These tests were performed for all three pairwise comparisons between broiler-breeders, broilers, and layers.

Previous work has shown that MD associated condemnation rates historically varied across broad geographic area such as between states (26). We explored whether there was evidence of clustering in virus detection across the finer spatial scales found in our cross-sectional data. We did this by calculating distances and correlations in effect sizes between each pairwise farm location. We then used the ‘lm’ function to generate two models. The first was an intercept only model that functioned as a null model. The second was an intercept plus distance effect model, where distance was transformed by adding one and then taking the log_10_. The importance of distance was assessed by performing a likelihood ratio test.

### Analysis of the longitudinal data

To study the variation in MDV dynamics within a focal chicken house over time, we used the quantitative values returned by qPCR analysis, rather than the presence-absence used for the cross-sectional data, because the quantitative data are more sensitive to changes in virus concentration. We assumed lognormal error in these quantities, because variation in qPCR data tends to occur on a log scale (40). In our analyses, we therefore transformed the virus-copy-number-per-mg-of-dust data by adding one and log_10_ transforming that value. We explored the suitability of this lognormal assumption for our data in Supplement A.9. For samples with virus concentrations below our limit of detection, we performed our analyses while treating these data in two different ways, first as a value of zero virus copies representing virus absence, and second as a value of our limit of detection representing virus presence at an undetectable level. Our limit of detection was generally better than 100 virus copies per mg of dust (Supplement A.4), and so in practice, we used this quantity as our value in the latter case. For this analysis, all samples that had detectable virus below this quantity were treated identically to negative samples.

We sampled from five broiler farms at approximately weekly intervals. One of our main goals was to quantify how virus concentrations changed over the duration of a cohort, and across different cohorts, and so we began by merely plotting the data. A similar plot was generated for the air tube data. We then explored a cohort age effect by fitting smoothing splines to the raw data from each farm where the data are sorted by cohort age. Each spline was fit using the function ‘smooth.spline’. We used the option “nknots=4” for this function because this was the smallest number of knots that did not return an error. Very similar conclusions were obtained using any number of knots from four to nine. We explored seasonality in these data by subtracting cohort age effects from the raw data and plotting the residual virus concentration. We assessed the degree of correlation between houses within farms using the ‘cor’ function. We also examined autocorrelations within houses using the ‘acf’ function for data within each house.

## Results

### Cross-sectional data

Summary statistics characterizing the data used for our model comparisons are shown in fig. 2. Among all samples collected (combining cross-sectional and longitudinal data), wild-type MDV was detected at least once on 36 of the 104 farms (fig. 3). Virus was detected in 1 of 3 broiler-breeder operations, 4 of 5 broiler operations, and 3 of 5 egg layer operations. The fraction of samples in which virus was detectable varied substantially among farms with detectable virus, and less so between houses within a farm (fig. 3). Summary plots of virus prevalence as a function of production type, bird age, date of sample collection, and bird sex can be found in Supplement A.10. Note, however, that a visual inspection of patterns in these data could be misleading because of potential confounding with other covarying factors. We therefore used statistical models to further explore the effects of these factors on the data.

**Figure 3:**
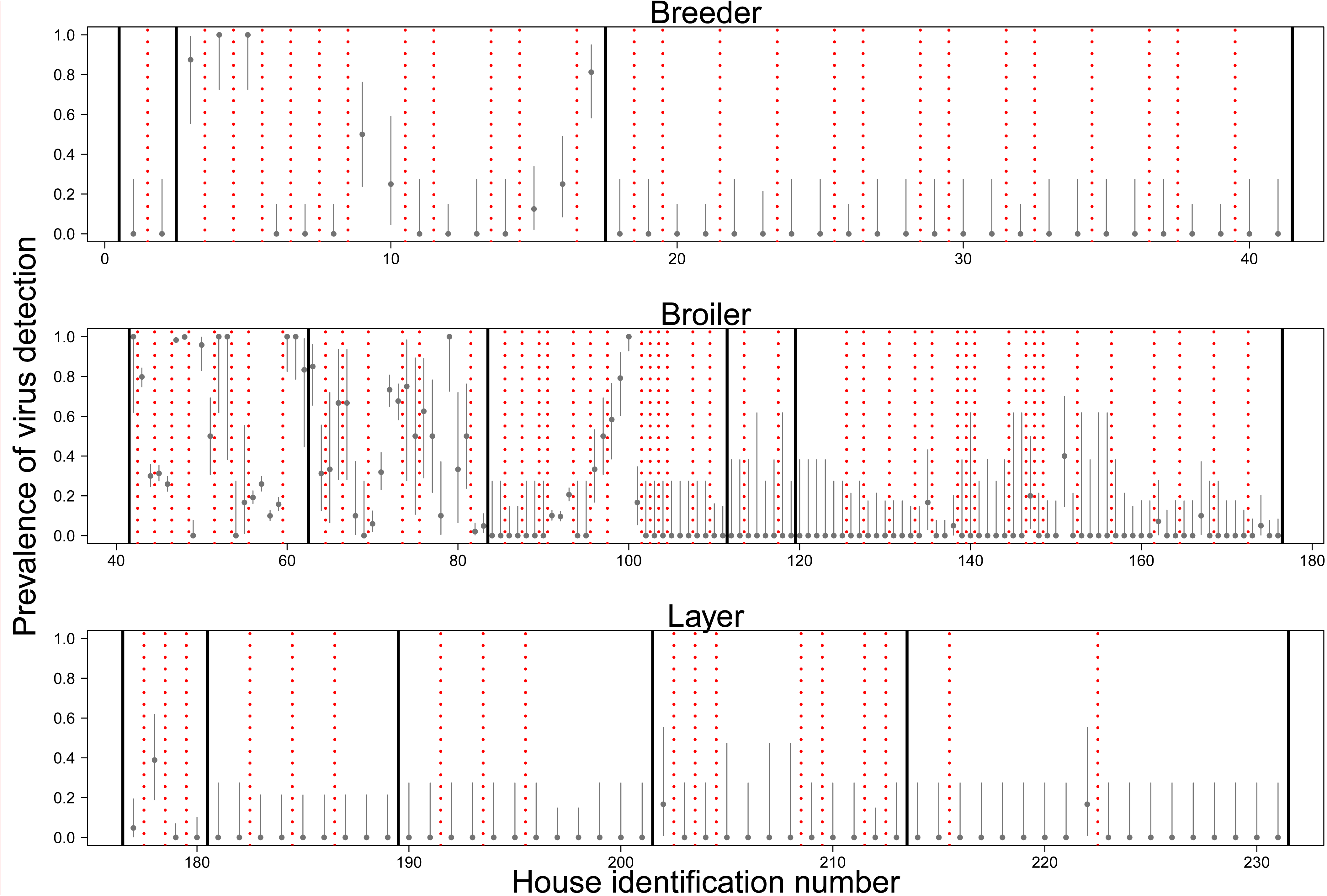
Fraction of tests with detectable virus. Each point shows the mean for a different house with grey bars depicting 95% confidence intervals on the mean (Supplement A.15). Confidence intervals vary between houses because of variable sample sizes. Different rows depict different production types (top–breeders, middle–broilers, bottom–layers). Solid black lines separate different operations (companies). Dashed red lines separate different farms. Note that prevalence estimates are from the raw data, not corrected to account for potential confounding effects such as bird age, collection date, or flock.

Our analysis of the virus prevalence data using DIC scores revealed that our best model was our most complicated model, which included effects of production type, bird age, collection date, and variation between dust samples (Table 1). Comparing our most complicated model to the other models through ∆DIC, we found moderate support for an effect of production type, reasonable support for an effect of collection date, relatively strong support for an effect of bird age, and overwhelming support for variation between dust samples. Taken together these results suggest that, to varying degrees, each of these factors had detectable effects on the prevalence of MDV on farms.

**Table 1:** Deviance information criterion (DIC) table for models considered. “Mean deviance” is the average deviance of the posterior. ∆DIC is defined as the difference in DIC between the model with the smallest DIC and the focal model. Note that the “Full model” is in bold to highlight that it was the best model according to DIC.

| **Model name** | **Mean deviance** | **Number of parameters** | **DIC** | **ΔDIC** |
| --- | --- | --- | --- | --- |
| **Full model** | **336.9** | **17** | **494.5** | **0** |
| No production type | 339.7 | 15 | 497.1 | 2.5 |
| No bird age | 345.8 | 15 | 503.7 | 9.1 |
| No collection date | 341.2 | 10 | 499.1 | 4.6 |
| No sample | 450.1 | 16 | 575.3 | 80.7 |

We further explored the importance of these effects by examining the fraction of variance in our data explained by each model factor for our best model (fig. 4). This showed that the fraction of variance attributable to production type was highly uncertain, with the 95% credible interval ranging from 1.5% to 38.4%.

**Figure 4:**
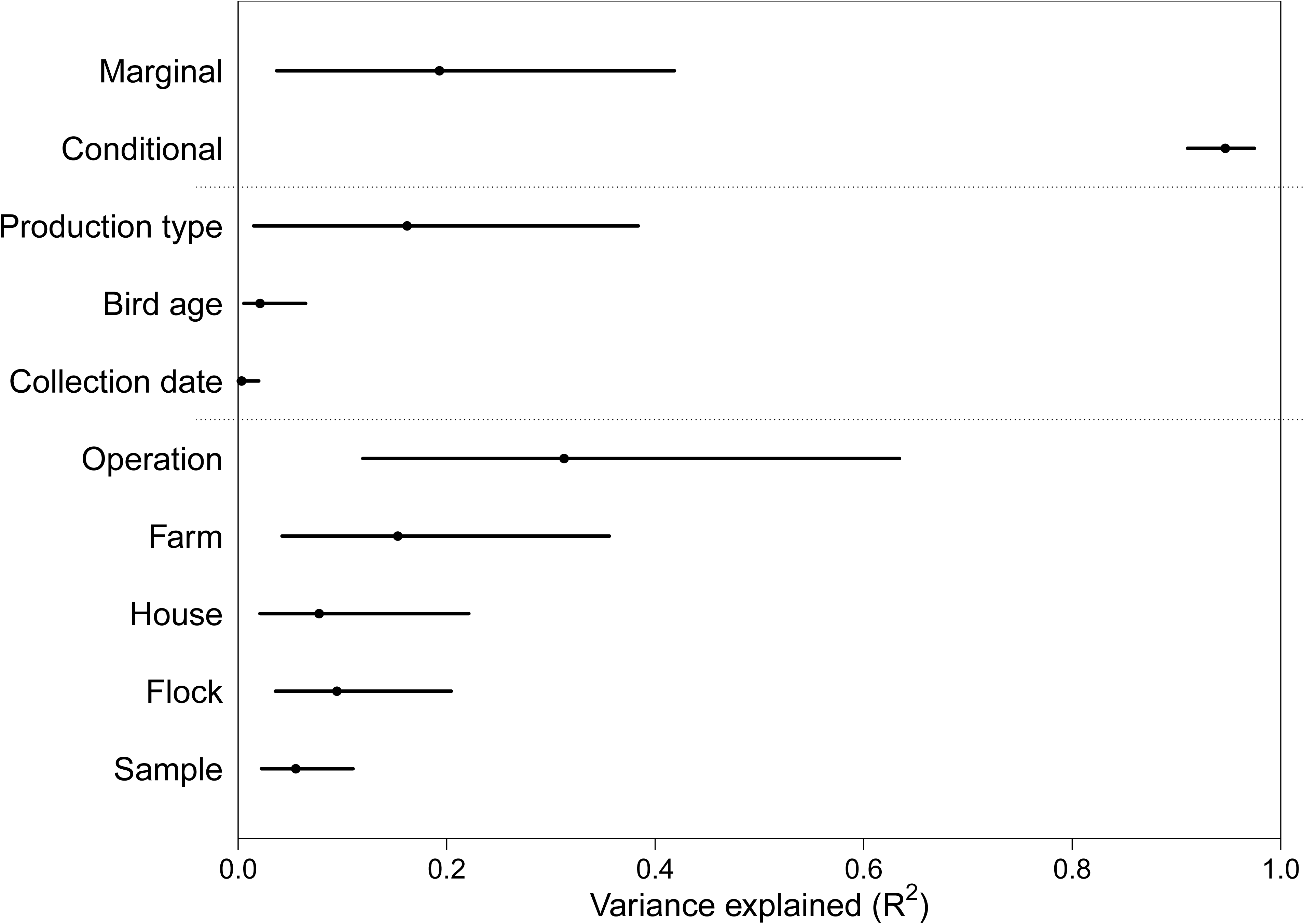
Fraction of variance on the latent scale attributable to each model factor. Points are median values and lines are 95% credible intervals. Marginal and conditional *R*^2^ values represent the variance explainable by all fixed effects, and all fixed plus random effects respectively. Note that only the values for the best model (Table 1) are shown.

The effect sizes of production type, bird age, and collection date observed in the full model are shown in fig. 5. Virus prevalence was higher on broiler farms than on layer farms (*p* = 0.02), but there was no statistically significant difference between breeder and broiler (*p* = 0.27), or breeder and layer farms (*p* = 0.15). During the first few weeks of a bird cohort the probability of detecting virus decreased, and then as birds continued to age this probability began to increase. Note that after cohorts reached about 100 days, the median effect was close to neutral and the confidence intervals on the effect size were fairly large (fig. 5 middle panel). This uncertainty was likely because we have relatively few data from older cohorts. We additionally saw a seasonal pattern in MDV prevalence with a fairly wide credible interval. Our probability of detecting virus was lowest in the winter months and highest in the summer months (fig. 5 bottom panel).

**Figure 5:**
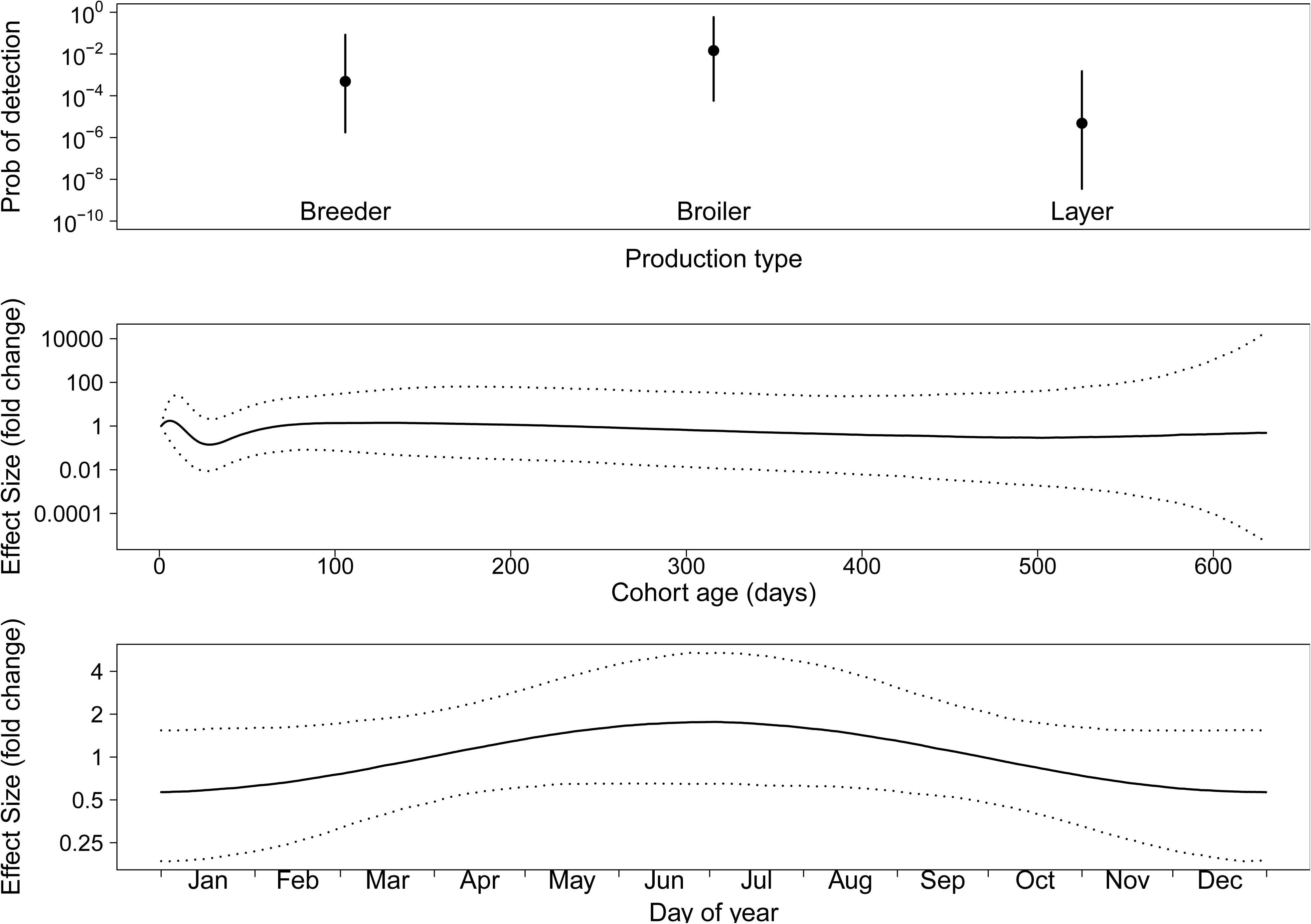
Effect sizes for fixed effects. The top panel shows the median and 95% credible interval for the three production types. The middle panel shows the median and 95% credible interval for the effect of bird age on the probability of detecting virus in a dust sample. The bottom panel shows the median and 95% credible interval for the effect of collection date on the probability of detecting virus.

Additionally, we found that the estimated effect that “Farm” had on virus detection tended to be positively correlated for nearby farms, and this correlation decayed with distance between farms (χ^2^ = 28.5, *d.f.* = 1, *p* < 0.001). However, the effect size was relatively small, with a maximum estimated correlation of 0.029 ± 0.004 that decayed by 0.014 ± 0.003 with every log_10_ increase in distance. Moreover, this correlation with distance might have been a statistical artifact resulting from geographic clustering of farms belonging to the same operation: no significant correlations by distance were detected between farms within single operations.

### Longitudinal data

The longitudinal data from five broiler farms revealed several patterns. These data visually confirmed the conclusion from the cross-sectional data that virus densities varied substantially between farms, and between flocks, but varied less between houses located on the same farm (figs. 6 and 7). This similarity between houses was also seen as a correlation of virus quantities between houses within farms (average correlations between houses within each of the five farms were 0.215, 0.320, 0.738, 0.763, and 0.918). The data also confirmed the observation that virus densities tended to decrease during the early phase of a cohort, and tended to increase during the later phase of a cohort (Supplement A.11). This created “U” shaped curves in virus concentration within cohorts (figs. 6 and 7). This pattern is not explained by differences in sample humidity or qPCR inhibition (Supplement A.12). Consistent with the cross-sectional data in which seasonal effects were small, we were unable to find any consistent seasonal effect on MDV dynamics in these data.

**Figure 6:**
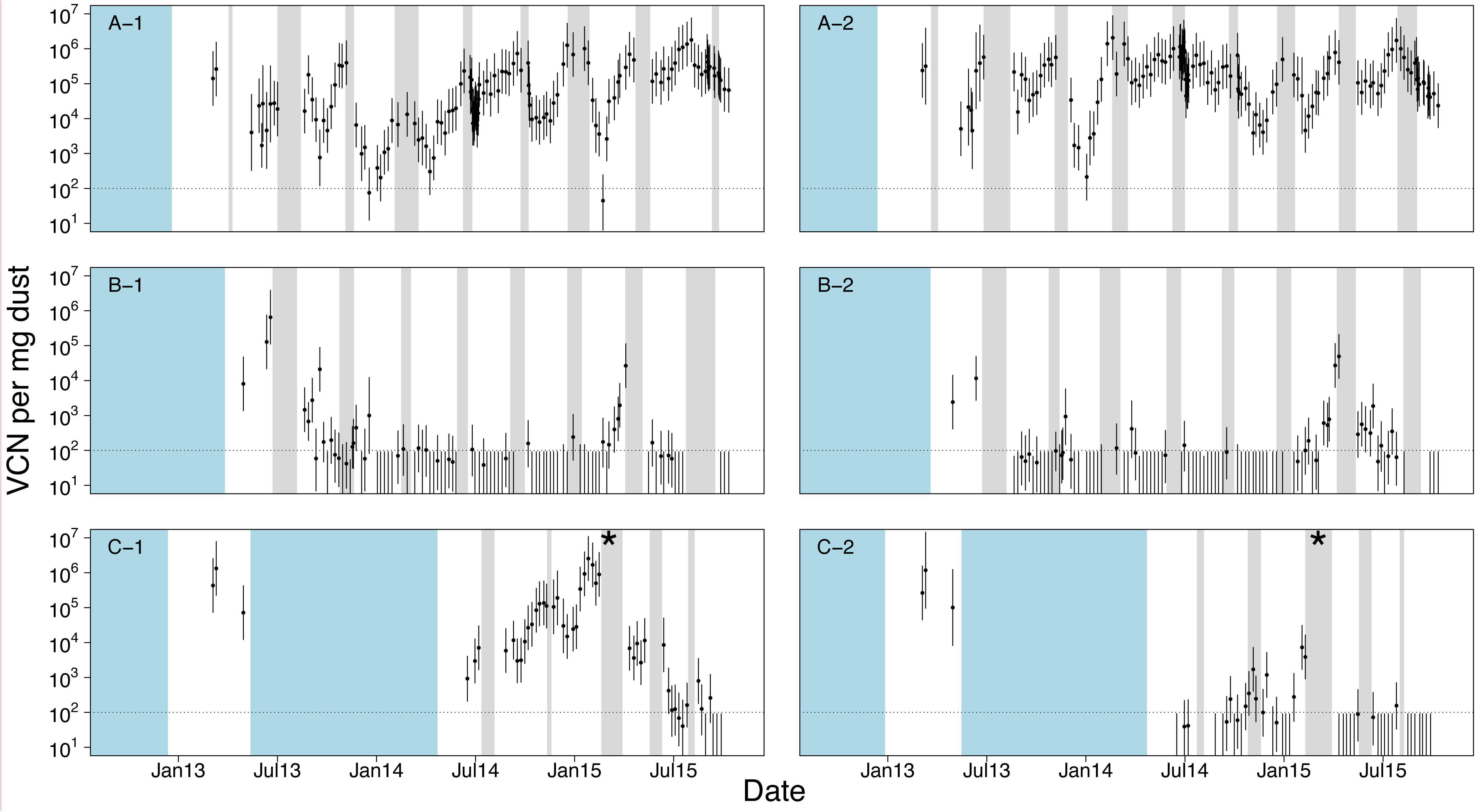
Longitudinal surveillance data for three broiler farms in Pennsylvania. Each panel is labelled “X-Y”, where “X” gives a unique farm identification, and “Y” gives a house number on that farm such that each two character label is unique. Each of the three farms shown in this figure had two houses. All of these farms began associated with the same operation, but farm “C” changed operations in the middle of our surveillance. The timing of this change is denoted by an asterisk in the plot. All farms followed an “all-in, all-out” policy meaning that houses had discrete periods of rearing and down time. To represent the presence or absence of birds, white intervals cover periods when birds were present, grey intervals cover periods when birds were absent, and blue intervals cover unknown periods. Each point represents the log-mean virus concentration (VCN) for that set of dust samples. Error bars are 95% confidence intervals calculated as explained in Supplement A.15. The dotted horizontal line shows the approximate qPCR limit of detection for a single test.

**Figure 7:**
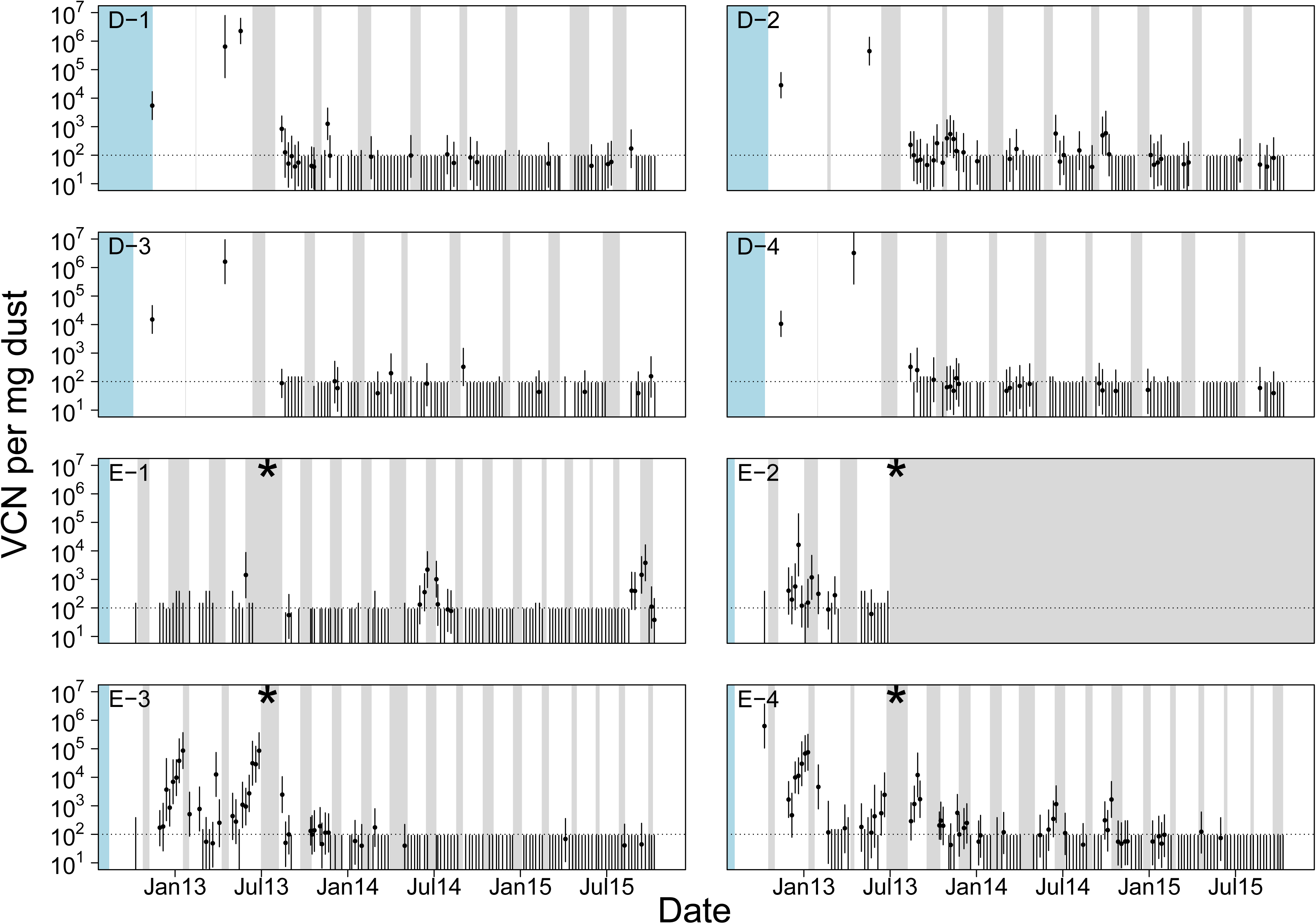
Longitudinal surveillance data for two additional broiler farms in Pennsylvania. Symbols, colors and layout as in fig. 6. Both of these farms had four houses. Farm “D” was associated with the same operation as the farms in fig. 6, but farm “E” was not. Note also that farm “E” changed operations during our surveillance period, the timing of which is marked with an asterisk.

Three additional patterns were also detectable in the longitudinal data. First, virus concentrations often dropped to below detectable levels, and returned to detectable levels at a later time point (figs. 6 and 7). Second, there was an autocorrelation in virus concentration within single houses over time. This effect was seen as an autocorrelation between samples collected seven days apart (Acf(7)_avg_ = 0.579, Acf(7) _min_ = 0.226, Acf(7) _max_ = 0.967), although this correlation was also observed over longer periods (Supplement A.13). Third, during farm down time, when birds were absent from houses, there were many cases where virus concentration did not change (figs. 6 and 7). Patterns consistent with the first two of these observations were also seen in the air tube and feather tip data (fig. 8).

**Figure 8:**
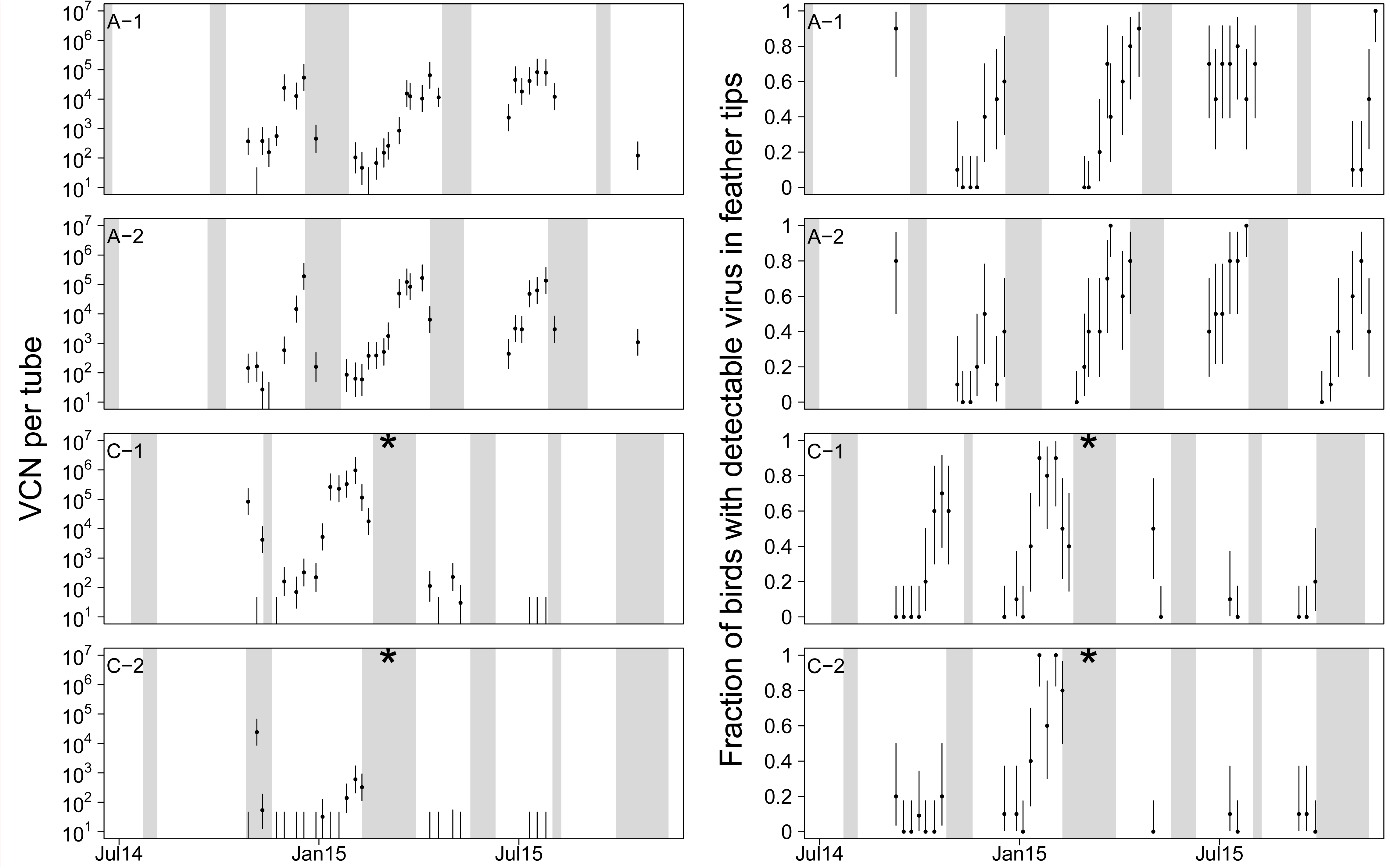
Air tube data (left column) and feather tip data (right column) for two broiler farms in Pennsylvania. Symbols, colors and layout as in fig. 6. Note that the dynamics in the air tube data and feather tip data are highly similar to one another, and are highly similar to that of the corresponding houses in the cross-sectional data (fig. 6). As in fig. 6, a change in operation on farm C is denoted by an asterisk.

## Discussion

We surveyed commercial chicken farms in Pennsylvania to generate the first industry-wide dataset exploring the prevalence of this virus in modern commercial settings. We found that the virus was detectable on only one third of farms, that bird age, collection date, and production type affected the probability that we detected virus, and that the vast majority of variation in the data was not attributable to those factors, but instead was attributable to differences between the companies, farms, houses, flocks and samples. Longitudinal sampling on five focal broiler farms revealed substantial autocorrelation in virus density within houses over time, and demonstrated that virus concentrations often dropped to undetectable levels on farms but reappeared in future flocks. Taken together, these results show that the virus can be found throughout the heterogeneity of the poultry and egg industry.

Despite the differences in rearing practices between the United States, Australia and Iraq, the overall prevalence of MDV detection in dust samples was broadly in agreement with studies performed in these other countries (17, 37, 43, 44) showing virus on only a subset of farms. Like Walkden-Brown et al. (44), we found that MDV concentration in dust increased in broiler flocks as birds aged. Two Australian studies examined the link between HVT and MDV concentration in dust. One study found no correlation (17) and the other showed a negative correlation (44). Our results agreed with the former study. All flocks in our study, however, were vaccinated, limiting the variation in vaccination status of our study relative to the studies performed in Australia where vaccination is not universal. One striking difference between our conclusion and that of Groves et al. (17), was our finding that operations have vastly different levels of MDV prevalence. Groves et al. (17) found no effect of operation. It may be that the importance of operation is specific to poultry farming in the United States.

Previous studies on the evolution of MDV in the poultry industry have focused entirely on endemic virus persistence in broiler chicken houses (1, 39, 41). Our data, however, reveal that the virus can be found in each of the sectors of chicken farming, including broiler, layer, and breeder chicken facilities. The assumption of these models, that virus evolution can be understood using the host genetics, rearing duration, host densities, vaccination strategies, and biosecurity measures employed in the rearing of broiler chickens alone therefore might be misleading. Given the potential for vastly different evolutionary outcomes under different ecological assumption, further investigation is needed to determine where evolution is likely strongest.

Conventional wisdom is that MDV is sufficiently pervasive that it should be considered ubiquitous (14, 30, 33). This idea came from observations that the virus is highly stable in the environment (24), that problems with MD can occur quickly and without warning when there are issues with vaccine administration, and that vaccination does not preclude infection with and transmission of the virus (22, 35, 38). It was further supported by the historical ubiquity of antibody detection in production flocks (5, 11, 20, 47). However, we found virus on only one third of farms. It may in fact be present on the other two thirds of farms at densities below our detection threshold or at times when samples were not collected, or it may instead be that modern farm practices have led to changes in the distribution of the virus such that it is no longer ubiquitous on chicken farms. Many features of poultry farming have changed in recent decades that could have altered MDV ecology, such as vaccination strategies and cohort durations (26, 41). Recent studies in Australia (37, 44), and Ethiopia (4) have suggested that MDV may no longer be ubiquitous in those locations. Our study suggests that this trend may be more general, extending to commercial poultry farming the United States. Introducing non-vaccinated sentinel birds could be a way to directly challenge this finding. If confirmed, this suggests that selective forces acting during sporadic outbreaks or acting in flocks with low prevalence of infection may play an important role in the evolution of the virus.

The importance of random effects (i.e. operation, farm, house, flock, and sample) in explaining the data suggests that substantial variation in virus dynamics are explained by factors that co-vary with these random effects. For example, bird breeds, vaccination details, and average cohort durations may explain some of the variation between operations. Ventilation rates, clean out efficiency, and other hygiene factors may explain some of the variation between farms. Structural differences and wind patterns may explain some of the variation between houses. Microbial communities, developmental plasticity and stochastic effects of virus transmission may explain some of the variation between flocks. Lastly, spatial clustering of virus may explain some of the variation between samples. Our model analysis showed that between about one quarter and three quarters of the variation in MDV detection probability was attributable to the combined effect of production type and operation. However, we are unable to parse these effects into more specific factors such as hygiene, barn design, ventilation, temperature, or vaccine manufacturers. This is because these factors strongly covary with factors such as production type and operation. For example, all layer and broiler-breeder farms used Rispens vaccination, and almost all broiler farms used bivalent vaccination. Nevertheless, our results suggest that factors outside the control of individual farm operators may play a large role in MDV dynamics. It also suggests that any intervention strategy intended to control virus is likely to be ineffective unless implemented through changes in operation practices or policies.

The large degree of uncertainty in the effect sizes of production type and operation likely resulted from correlations in these estimates (Supplement A.14), and this correlation may explain why support for an effect of production type was only moderate. Indeed, exploring the variance explained by these two factors combined, we found that they accounted for between 26.7% and 74.4% of the variance. This parameter estimation difficulty likely occurred because these factors covary in our study area.

The observation that seasonality explained only a small portion of variance in MDV prevalence contrasts with observations that MD associated condemnation in broiler chickens has had clear seasonal patterns in the past (44, 45). However, seasonal patterns in condemnation have become less pronounced in recent years (26). The data we report here are consistent with the theory that this decrease in seasonality is attributable to an overall decline in prevalence, resulting in stochastic outbreaks playing a relatively larger role in dynamics than seasonal forcing (26).

The “U” shaped pattern in virus dynamics within a flock, seen both in the longitudinal and cross section data, suggests that MDV density in dust changes predictably over time. The initial decrease might be explained either by a dilution of virus in dust early in cohorts when birds shed virus-free dust into dust that remained from the previous cohort, or by degradation of virus DNA early in flocks. The subsequent increase could then be explained by the hyper-concentration of virus in dust as cohorts aged, when birds were shedding dust that was highly contaminated with virus.

In this study, the majority of data were collected from dust samples scraped from surfaces. An alternative method would have been the use of settle plates that collect dust as it settles out of the air. Both methods introduce biases, but we opted for the former method to avoid spatial artifacts that might have arisen from patterns of dust flow. Our measurements of virus concentration showed little evidence of spatial heterogeneity (Supplement A.2). Perhaps the largest drawback of our method was that each sample of dust potentially contained material that might even predate the current flock of birds in the house. The dust kinetics might therefore be dampened relative to their true kinetics in the air. However, the strong agreement in viral kinetics between these data, and both the air tube and feather tip data suggest that this is may be more of a theoretical rather than practical concern.

An interesting question is whether virus populations are persisting within individual houses and farms, or instead going through repeated extinction and recolonization events. Our observation in the longitudinal data that there was a strong autocorrelation in virus concentration within houses over time (Supplement A.13) contrasted with the observation that virus densities were often undetectably low in one cohort but emerged as detectable in the next (figs. 6 and 7). This reemergence might be due either to recolonization events or to the epidemiological amplification of virus persisting within the house at undetectable concentrations. Recently developed genetic sequencing techniques (32) could be used to determine the relative contributions of these two factors.

## Acknowledgements

We thank the participating poultry companies of Pennsylvania for providing access to farms and to flock information. We thank John Dunn for supplying us with the prMd5pp38-1 plasmid. This work was funded by the Institute of General Medical Sciences (R01GM105244), National Institutes of Health and United Kingdom Biotechnology and Biological Sciences Research Council as part of the joint NSF-NIH-USDA Ecology and Evolution of Infectious Diseases program, and by the RAPIDD program of the Science and Technology Directorate, Department of Homeland Security and Fogarty International Center, National Institutes of Health (DAK, AFR). The funders had no role in study design, data collection and analysis, decision to publish, or preparation of the manuscript.

